# Visual mode switching learned through experience

**DOI:** 10.1101/2020.07.17.209478

**Authors:** Yanjun Li, Katherine EM. Tregillus, Qiongsha Luo, Stephen A. Engel

## Abstract

When the environment changes, vision adapts to maintain accurate perception. For repeatedly encountered environments, learning to switch immediately to prior adaptive states would be beneficial, but past work remains inconclusive. We tested if the visual system can learn such visual mode switching for a strongly tinted environment, where adaptation causes the dominant hue to fade over time. Eleven observers wore red glasses for five one-hour periods per day, for five days. Color adaptation was measured by asking observers to identify “unique yellow”, appearing neither reddish nor greenish. As expected, the world appeared less and less reddish during the one-hour periods of glasses wear. Critically, across days the world also appeared significantly less reddish immediately after donning the glasses. This indicates that the visual system learned to shift rapidly to a partially adapted state, switching modes to stabilize color vision. Mode switching likely provides a general strategy to optimize perceptual processes.

## Introduction

When the visual system encounters different environments -- for example a change in overall brightness, focus, or color – sensory processing also changes, in order to maintain accuracy and efficiency. Some of the processes producing such adjustments, called visual adaptation, unfold gradually (Clifford et al., 2007; Kohn, 2007; Wark et al., 2007; Webster, 2015). For example, wearing sunglasses can alter the color of an apple, making it difficult to determine if it is ripe. But as our visual system adapts, the apple’s apparent color gradually returns to normal. For common environmental changes, it would be beneficial if the visual system could remember past adaptation, and rapidly switch to the appropriate state (Engel et al., 2016; Yehezkel et al., 2010). Such visual mode switching would aid the many functions that adaptation serves, including improving the detection or discrimination of objects and their properties (Dragoi et al., 2002; Krekelberg et al., 2006; McDermott et al., 2010; Müller et al., 1999; Wissig et al., 2013) and making neural codes more efficient (Seriès et al., 2009; Sharpee et al., 2006; Wainwright, 1999).

Empirical evidence for learning to switch rapidly to an adapted state is sparse and inconclusive, however. A few studies have found preliminary support for learning effects on visual adaptation (Engel et al., 2016; Yehezkel et al., 2010), but others have found little to no effect of experience (Tregillus et al., 2016; Vinas et al., 2012). Notably, previous work has not measured the effects of moving in and out of an environment multiple times per day over many days, and none has tested for changes in the time course of adaptation with experience. Thus, it remains unclear whether people can learn to rapidly switch visual modes with experience.

Here, we used color adaptation to test for such learning: Observers wore a pair of tinted glasses, which made the world appear very reddish (the spectral transmission of the glasses as well as the monitor gamut with and without the glasses are shown in Fig. 1). Color adaptation in such situations is relatively well-understood, and one of its main effects is that the dominant color of the environment fades over time e.g. (Belmore & Shevell, 2008; de La Hire, 1694; Eisner & Enoch, 1982; Neitz et al., 2002; von Kries, 1902), restoring the world to its prior, “normal” appearance.

**Fig. 1.**
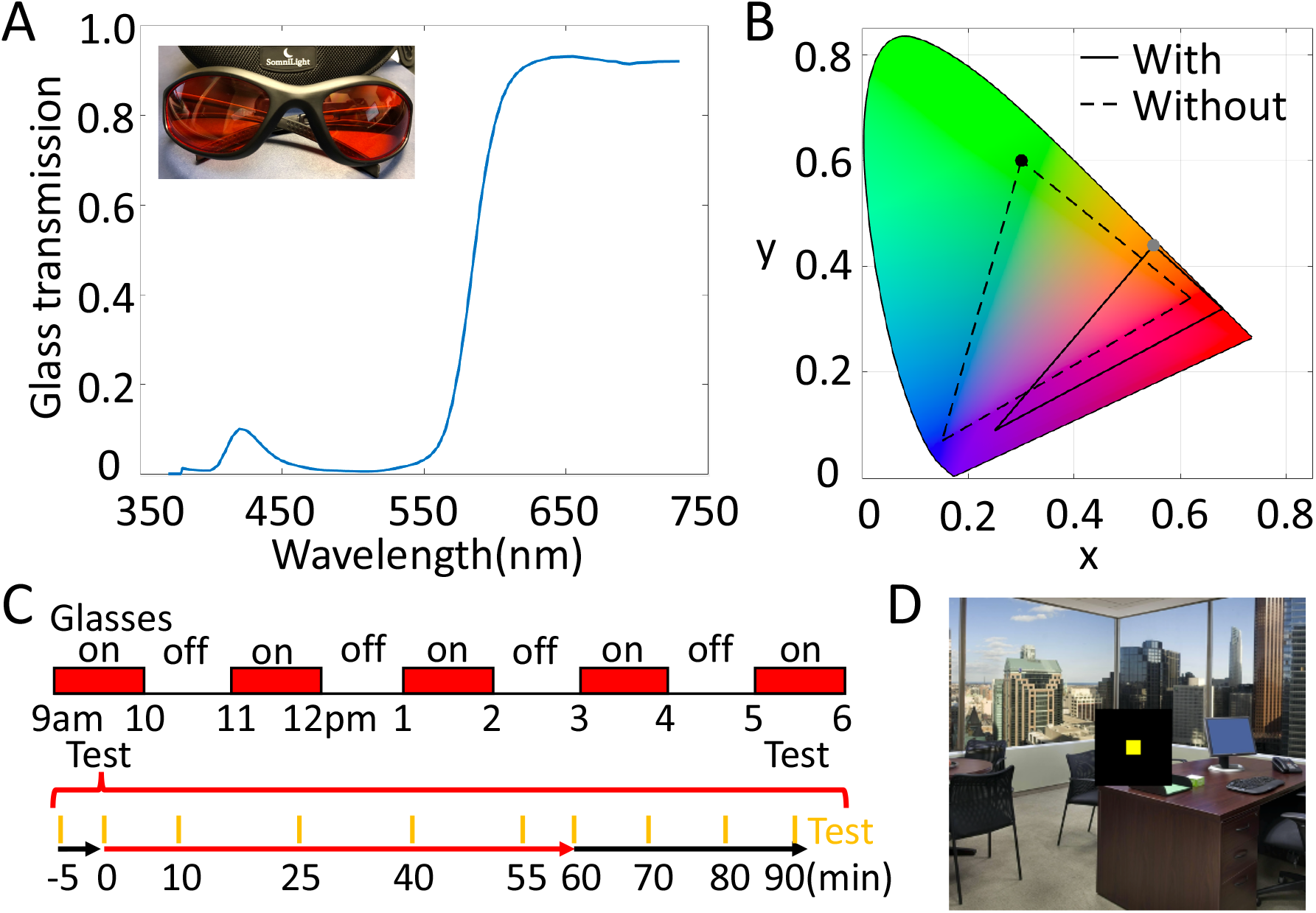
Glasses transmission and experimental procedure. **A)** The red glasses used in this study and their transmission spectrum. The glasses filter out most of the energy at short wavelengths and maintain most of the energy at long wavelengths. **B)** Monitor gamut with (solid line) and without (dashed line) the glasses plotted in CIE color space. The glasses compress the gamut and shift it towards red chromaticity. For example, the greenest light produced by the monitor (black dot) falls in an orange part of color space through the red glasses (gray dot). **C)** Experimental procedures. The upper panel indicates the times when the observers wore the glasses within one day. Two test sessions were conducted, during the first and last 1hr of wearing the glasses. The lower panel illustrates the test procedure in each session. Orange bars indicate the time of test: 5 min before putting on the glasses, right after putting on the glasses, then following 10 min, 25 min, 40 min, and 55 min of wearing the glasses. Observers then removed the glasses and were tested immediately, and 10 min, 20 min, and 30 min after removing the glasses. **D)** Test display. Observers adjusted the color of a square centered on a background image of a naturalistic environment, presented on a monitor in a fully lit room. A black square of 5.7 degrees separated the 0.5 deg square test patch from the background image. The test patch was presented for 200 ms at 1.5 sec intervals, and the observer’s goal was to set it to appear unique yellow.

Observers in the present experiment donned and removed the glasses multiple times a day for 5 consecutive days. We hypothesized that rapid color adaptation would strengthen over days, such that observers would experience a much smaller perceptual change in the color of the world when they put on the red glasses, providing evidence that they had learned to switch visual modes, to regain a prior adaptive state.

Observers wore the red glasses for five one-hour periods, each separated by one hour without glasses (Fig. 1). To track adaptation, we asked observers to make unique yellow settings, identifying the wavelength of light that appears neither reddish nor greenish. Unique yellow is a commonly used measure in color perception, in part because observers are highly consistent in their judgments. On each day, observers were tested in two sessions, once in the morning and once in the afternoon, for 5 days in a row. In each test, observers made unique yellow settings for 5 minutes. In each session they performed one test before putting on the glasses; 5 with the glasses on; and 4 after removing the glasses. The tests were all conducted in a fully lit room in order to provide information about the visual environment present. In a follow-up, conducted about one month after the main experiment, observers participated in one additional and identical testing session.

## Results

The world appeared very reddish when observers first put on the glasses, and the redness faded over time as vision adapted. Fig. 2 plots the unique yellow settings (quantified as hue angle, see Methods) as a function of time, averaging across 11 observers, for the 5 days. The relatively small number (around 220) on the very first test with glasses on (red dots) indicates that observers’ unique yellow was physically relatively green, which was required to cancel the redness produced by the glasses. The upward slope of each session’s 5 settings shows that observers added less green to unique yellow over time, adapting to the red environment during the 1hr of wearing the glasses, with the world looking less and less red. This pattern can be seen both in the morning (with white background in Fig. 2) and the afternoon (with light gray background) session on all 5 days.

**Fig. 2.**
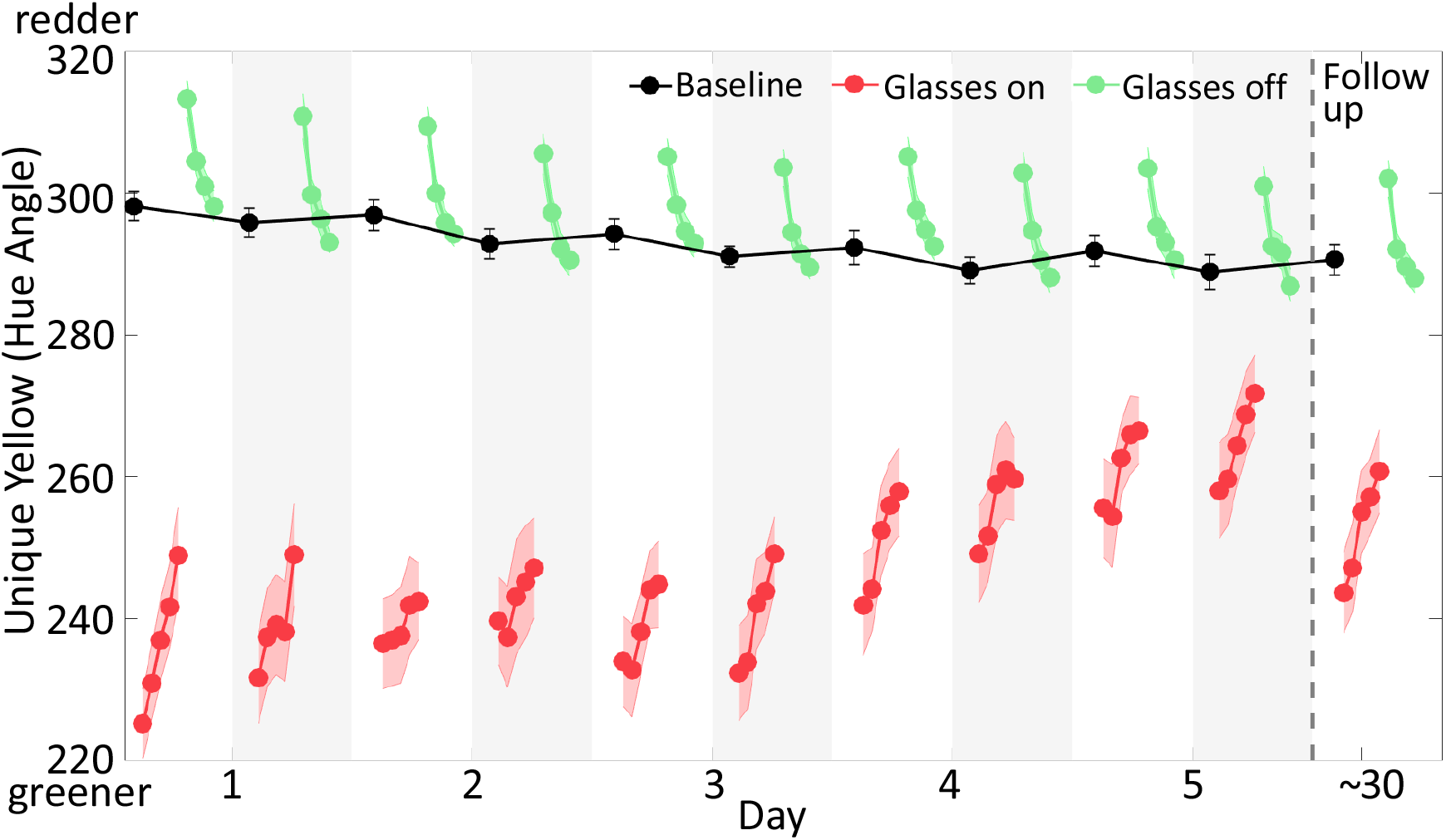
Results of the main experiment and the follow-up tests. Mean unique yellow settings represented in hue angle are plotted as a function of time for 5 days and the follow-up test. The black dots are baseline settings, made at the beginning of each test session with glasses off. The white background indicates morning sessions, and the light gray background indicates afternoon. The red dots plot settings with glasses on and the green dots are settings after removing the glasses. Successive symbols are plotted for each 5 min test (see Fig 1C). The red and green bands represent standard errors of the mean, computed across participants (N = 11).

### Rapid adaptation to the glasses strengthened across days

Across days, observers learned to rapidly adjust to the red glasses. That is, when they first put the glasses on, the world appeared less and less reddish. This is visible in the graph by the rising trend of the first unique yellow setting in each session across days. A linear trend analysis (Fig. 3 red dots and black dashed line) showed that this increase was reliable (*y*_t_ = 4.06t − 76.7 + *e*_t_, t = 6.87, *p* < 0.0001). We interpret these changes as increases in the amount of adaptation the visual system produced immediately upon putting on the glasses.

**Fig. 3.**
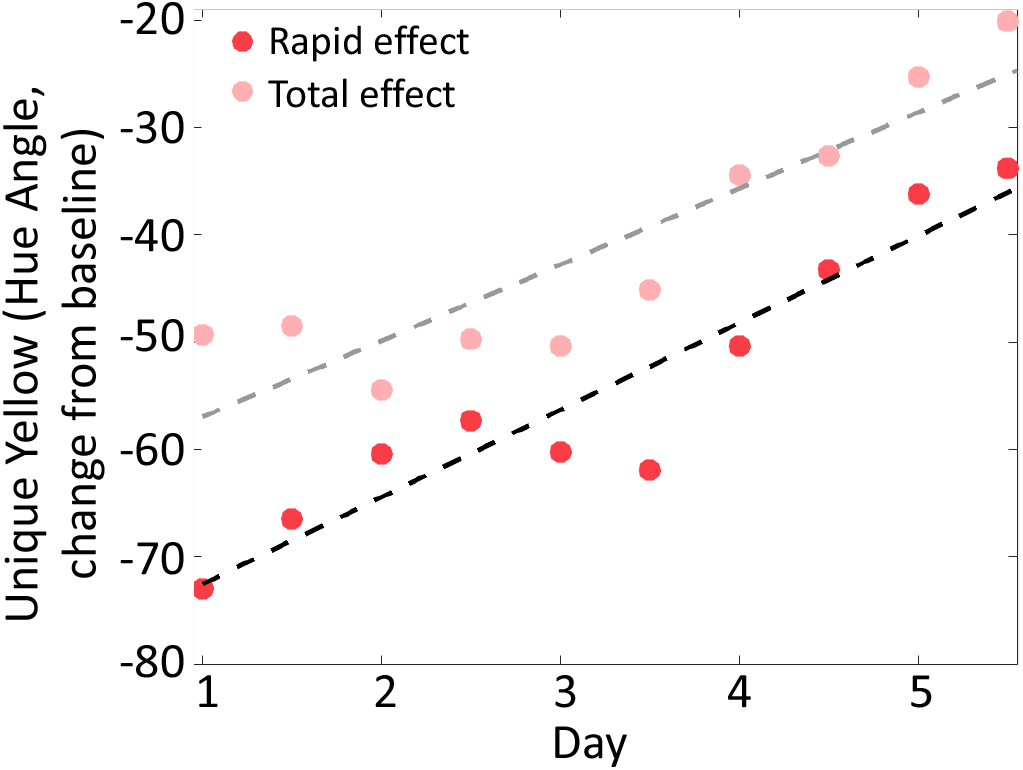
Rapid and total adaptation effects on 5 days. Red dots show rapid effects of adaptation, which are mean settings from the first 5 min test of each session with the glasses on. Total adaptation effects, denoted by the pink dots, are mean settings from the test taken after 1hr of wearing the glasses. Data have been corrected for possible baseline shifts by subtracting the baseline value for each session, taken immediately before putting the glasses on. The black and gray dashed lines are linear fits to the rapid and total adaptation effects, respectively. Both rapid and total adaptation effects grew significantly over days.

The changes were not due to lingering overall adaptation across days, as baseline measurements made before putting the glasses on showed a very different trend (see below).

How rapidly did this adaptation arise? Each data point in Fig. 2,3 represents average unique yellow settings from five one-minute blocks of testing. To better judge the timing of effects, we repeated our analysis using observers’ averaged settings within only the first one-minute block. We also repeated the analysis using observers’ very first unique yellow setting in each initial block, which happened within 20 seconds. In both cases, the first point after donning the glasses again shifted significantly across days (p < 0.001 for the first block, p < 0.03 for the first setting), suggesting that the observers achieved a relatively rapid switch between adaptive states (Fig. S2).

The amount of gradual adaptation to the red glasses during the 1 hour of testing, on the other hand, did not change across days. To estimate the amount of gradual adaptation within the 1 hour, we calculated the slope of the unique yellow settings within each test session. The grand average slope was 13.30 degrees of hue angle towards red in 1 hour, and there were no significant changes in slopes across test sessions (ANOVA, F_9,100_ = 1.06, p = 0.40). Given the increasing rapid and constant gradual effects, total adaptation, i.e. the sum of the rapid and the gradual adaptation, quantified by the last setting with glasses on in each session, increased across days (*y*_t_ = 3.95t − 69.6 + *e*_t_, t = 6.13, p <0.0001, Fig. 3 pink dots and gray dashed line).

### Learned switching between adaptive states was long-lasting

About one month (36±7 days) after the main experiment, observers returned for a follow-up test. (Fig. 2, right). Rapid adaptation to the glasses remained strong; the first block of unique yellow settings was redder than the settings from the first day of the main experiment (t = −4.83, p < 0.001). However, the effect was somewhat diminished, as the follow-up settings were greener than those made on day 5 of the main experiment (t = 3.28, p < 0.01). About 66% of the change across the 5 days was maintained in the follow-up test.

### Color aftereffects did not change across days

When observers removed the red glasses, they experienced a classical color aftereffect (Helmholtz, 1924; Krauskopf & Karl, 1992; van Lier et al., 2009), and reported the world looked slightly greenish, thus they added red to cancel out this aftereffect when making their unique yellow settings (Fig. 2, green dots). We tracked the decay of the aftereffect for half an hour after removing the glasses, as observers’ settings shifted back towards baseline. The decay followed a roughly exponential shape, as previously reported for color aftereffects (Fairchild & Lennie, 1992; Fairchild & Reniff, 1995; Wright & Parsons, 1934). The rate of the decay, as measured by an exponential fit, did not change over days (F_9,96_ = 0.01, p = 1).

### Baseline unique yellow became slightly greener across days

Baseline values of unique yellow on each day were measured as the mean setting from the first 5-minute test of the morning session, made before putting the glasses on; these settings were preceded by many hours (averaging approximately 15) since the glasses were last worn and were made without the glasses on. We observed a small but significant shift in baseline unique yellow settings over time, visible in Fig. 2 (black dots) as the hue angle baseline shifted towards green chromaticity (t = −3.33, p < 0.01). This is surprising because adapting to the red glasses makes redness more neutral over time, thus resulting in redder unique yellow (see Discussion).

To make sure our main finding of greater rapid adaptation did not depend upon this shift in baseline, we corrected its effect by subtracting the baseline setting in the morning test session on each day from all settings within the day. These baseline-corrected results showed an identical overall pattern across days as the uncorrected data, though some became slightly larger (Fig. S1).

### Color constancy increased across days

Color constancy, an important benefit of adaptation, is the extent to which objects appear the same color despite changes in viewing conditions (Brainard & Radonjić, 2014; Foster, 2011; Witzel & Gegenfurtner, 2018). Such stability against transient features of the environment allows color appearance to provide reliable information about object identity and state (e.g. the ripeness of an apple). If observers in our experiment had perfect color constancy, then the same physical color on the monitor, that is the same spectrum of light leaving the display, should appear unique yellow to observers both with and without the glasses, despite the glasses’ dramatic effect on the spectrum of light reaching the eye.

To estimate the amount of constancy, we followed previous work (Brainard & Stockman, 2010) and characterized the physical color reaching the eye using the relative absorptions of the long-wavelength (L) and medium-wavelength (M) photoreceptors: L/(L+M). This index is convenient because empirically measured unique yellow in baseline conditions is close to a balance point of 0.5.

Fig. 4 plots our results in this space and shows that color constancy improved across days. The black dots are baseline unique yellow settings before putting on the glasses; as expected, they fell around 0.5. The red dashed line at the top of the plot reflects perfect color constancy with glasses on, calculated by assuming that the physical color corresponding to unique yellow did not change from baseline on the first day. This identical spectrum of light would of course result in very different cone absorptions with the glasses on than off, because of the glasses’ effect on the light reaching the photoreceptors. On the other hand, if observers completely lacked color constancy, unique yellow settings with glasses on would simply remain at baseline values.

**Fig. 4.**
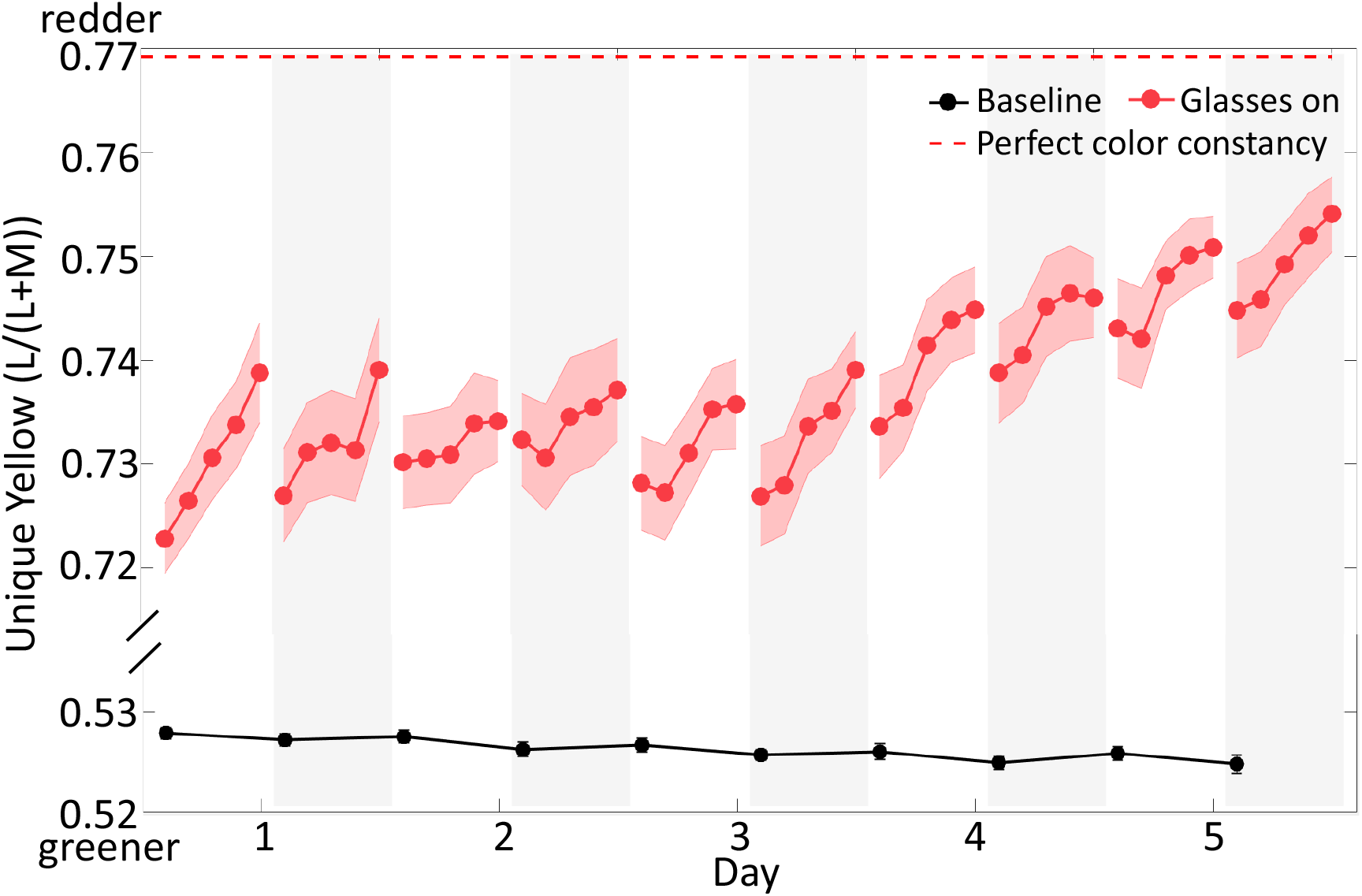
Mean unique yellow settings across 5 days, plotted as relative cone absorptions L/(L+M). The red symbols show the relative cone absorptions for settings with glasses on, corrected for the red glasses transmittance. The black dots are baseline settings taken at the beginning of each test session with glasses off. If the observers showed complete absence of color constancy, the unique yellow settings with glasses on should have been at the same level as this baseline. The red dashed line above corresponds to the baseline unique yellow corrected for the red glasses’ transmittance. If observers had perfect color constancy, their settings would produce identical physical colors on the monitor with and without glasses, and so should fall here when glasses were worn.

Across days, observers’ unique yellow settings (red dots) steadily rose towards the perfect color constancy line, indicating that color constancy improved. The very first time they put on the glasses, observers showed about 80% of perfect constancy, as calculated by the ratio between 1) the Euclidean distance between baseline and the first unique yellow setting with glasses on and 2) the distance between baseline and perfect constancy. This pre-existing constancy was presumably due to the rapid adaptation that produces the color constancy we experience in most situations (Rinner & Gegenfurtner, 2000; Smithson & Zaidi, 2004; Webster & Mollon, 1995). The amount of constancy grew significantly as observers learned to immediately switch to the appropriate adaptive state (t = 4.60, p < 0.001), and exceeded 90% on the 5th day.

### Individual differences in learning

Fig. 5 plots individual differences in changes in adaptation across observers. Some observers showed a large increase in the amount of rapid adaptation over five days (upper panel in Fig 5A, gray circle in Fig 5B), while others demonstrated a flatter pattern (lower panel in Fig 5A, black circle in Fig 5B). To test if the individual differences were statistically significant, we computed the Pearson correlation between the changes in rapid adaptation from the first day to the fifth day, and from the first day to the follow-up test. This correlation was significant (r = 0.81, p = 0.003, Fig. 5B), indicating that observers who had a larger learning effect over 5 days also retained larger amounts a month later, a form of test-retest reliability. Thus, individuals differ in their ability to learn to rapidly switch between the two states.

**Fig. 5.**
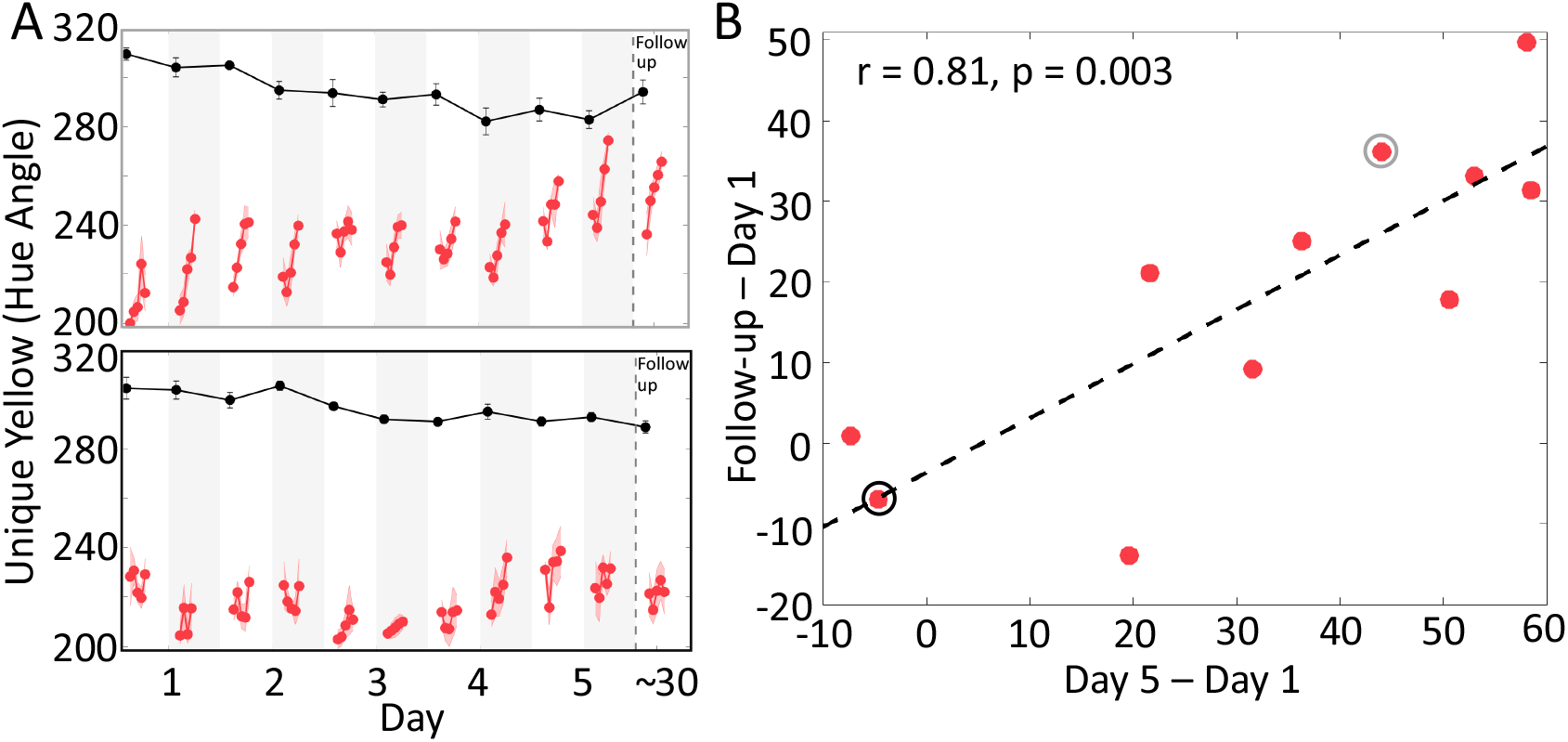
Individual differences in learning effect. **A)** Some of individual data. Results are from 2 observers in the study. One observer (upper panel) showed a gradual increase of rapid adaptation during the five days. This observer also retained the strong rapid adaptation in the follow-up test. Another observer (lower panel), showed a flatter pattern across days and little effect of learning in the follow-up test. **B)** Test-retest reliability of individual differences. The change in rapid adaptation to the glasses from the 1st day measured on the 5th day significantly correlated with the change measured in follow-up test, across observers. This indicates observers differed in their ability to learn to rapidly switch between the two states. Red dots represent observers and the dashed line is the least-square fit. The light gray and black circles denote the individuals plotted in the upper and lower portion of panel A, respectively.

## Discussion

Observers achieved stronger rapid adjustment to the red glasses through experience. Functionally, visual adaptation allows us to compensate for changes in the visual environment (Dragoi et al., 2002; Krekelberg et al., 2006; McDermott et al., 2010; Müller et al., 1999; Wissig et al., 2013) while also improving neural coding efficiency (Seriès et al., 2009; Sharpee et al., 2006; Wainwright, 1999). In situations where different visual environments alternate frequently, like wearing and removing glasses, the visual system repeatedly readjusts itself, to keep us seeing optimally in each. Our results suggest that observers can learn to switch between two adaptive states (i.e. to viewing with and without the red glasses) more efficiently over time. Such visual mode switching enables people to better handle the demands of the complex and changing visual environment.

Past work examining visual mode switching has produced mixed results. For example, observers who adapted to cylindrical lenses, creating a sort of astigmatism, showed fast re-adaptation in a second testing session (Yehezkel et al., 2010). However, clinically astigmatic observers showed little change in adaptation during 6 months following their initial prescription of corrective lenses (Vinas et al., 2012). Conflicting results also appeared in color perception, where in one study adapting to yellow filters produced little change in adaptation across 5 days (Tregillus et al., 2016), while another report showed that long-term habitual wearers of red and green lenses can adapt more rapidly than naïve observers to the color changes the lenses produce (Engel et al., 2016). Variability in observer populations and experimental procedures may account for these mixed findings. Our paradigm differed from past work in that observers adapted to very strong perceptual changes multiple times a day, and we tracked the detailed time course of adaptation in a test setting with rich cues to context (see below). Together, these factors likely produced larger changes and more reliable measurements of adaptation than observed previously.

Past work on long-term adaptation to colored environments, e.g. wearing red glasses or living under red lights continuously for part of the day, has found that adaptation grows stronger over days (Belmore & Shevell, 2008, 2011; Eisner & Enoch, 1982; Hill & Stevenson, 1976; Kohler, 1963; Neitz et al., 2002). However, these studies did not measure the time course of adaptation, or if observers could learn to rapidly switch between the different viewing conditions.

These past results were also highly variable, both within and between studies (Belmore & Shevell, 2008, 2011; Eisner & Enoch, 1982; Eskew & Richters, 2008; Hill & Stevenson, 1976; Kohler, 1963; Neitz et al., 2002; Tregillus et al., 2016), similar to the inconsistency in results on learning of adaptation. One reason for this variability may be that observers were tested with little context present, which may have made it difficult for the visual system to identify which environment it was in. For example, most tests were made in a completely darkened room, presenting only a single small test patch, making it difficult for the visual system to determine what adaptive state it should be in. The test setting in our experiment provided many cues that the visual system could use to tell which environment was present, i.e. whether the red glasses were on or off. These context cues may be necessary for mode switching to occur, though precisely which cues are important for which environments remains to be determined.

Unexpectedly, we found that the baseline unique yellow setting immediately prior to the introduction of the red glasses shifted towards physically more greenish across days. The shift was in the opposite direction from the color that the glasses produced and from the shift of the adaptation effect within 1hr. A similar trend in baseline settings was also found in two previous studies (Engel et al., 2016; Tregillus et al., 2016). While we can only speculate as to the cause of this pattern, it could be due to the aftereffect following the glasses’ removal. At that point, observers' judgments indicated that the world looked greenish to them, consistent with classical color aftereffects (Helmholtz, 1924; Krauskopf & Karl, 1992; van Lier et al., 2009). Adaptation across days to this greenish tint could have produced a shift in unique yellow towards green when not wearing the glasses. Long-term adaptation to aftereffects appears to be possible in other domains(Murch & Hirsch, 1972; Sheth & Shimojo, 2008).

The strengthened rapid adaptation we observed substantially improved observers’ color constancy, i.e. the stability of the perceived color despite the changes in viewing conditions (Brainard & Radonjić, 2014; Foster, 2011; Witzel & Gegenfurtner, 2018). Rapid adaptation, and even faster processes including 'simultaneous' local contrast, are likely major mechanisms that serve this constancy, e.g. (Rinner & Gegenfurtner, 2000; Smithson & Zaidi, 2004). A current debate in the field is whether constancy is improved for familiar, natural illuminant changes, which our visual systems may have encountered most often (Rüttiger et al., 1999; Delahunt & Brainard, 2004; Pearce et al., 2014; Radonjić & Brainard, 2016; Weiss et al., 2017). Our results suggest that training with repeated exposure can improve color constancy, at least for a very strong and unfamiliar illumination change. Observers show some amount of color constancy, and a variety of other perceptual constancies, in most natural settings, without any training. The extent to which these forms of visual mode switching are inborn, determined during development, or learned as an adult remains under investigation (Jameson & Hurvich, 1989; Sugita, 2004; Yang et al., 2015).

What mechanisms produced the faster switching between adaptive states? Neurally, adaptation to changes in the dominant color has effects on several sites within the retina (Boynton & Whitten, 1970; Lee et al., 1999; Rieke & Rudd, 2009) as well as cortical stages of color processing (Engel & Furmanski, 2001; Rinner & Gegenfurtner, 2000). One hint to the neural locus in our experiment is that changes were not observed in either adaptation within the hour of glasses wearing or in the size and decay of the color aftereffect when the glasses were removed. This independence from classical adaptation effects, which partly arise early in the visual system, suggests that mode switching may arise relatively late in processing (Rinner & Gegenfurtner, 2000). Identifying more precisely the extent to which learning can affect these different stages of processing could be profitably addressed in the future.

Computationally, one can view adaptation as the result of an inference process, in which the visual system must determine whether the visual environment has changed (Grzywacz & de Juan, 2003; Kording et al., 2007; Wark et al., 2009). Through exposure to the alternating colored and uncolored environment, observers in our experiment may have learned: 1) that the red environment was more likely (i.e., it had higher prior probability); 2) to more efficiently extract evidence of the red environment (giving it a higher likelihood); 3) that the red environment was likely to persist for a long time (making it costly to not adapt); 4) to speed inference by remembering, rather than re-inferring, the past adaptive state for the red environment. All these possibilities could produce stronger immediate adaptation, and they are not mutually exclusive. Future work could determine which factors are responsible for the changes of adaptation effects over time.

In sum, our results demonstrate that the visual system can learn to rapidly switch to prior adaptive states. This mode switching lessens the perceptual changes produced by changing viewing conditions, which could help in a number of perceptual tasks, for example recognition of objects or materials, discrimination between similar objects or materials, as well as improved communication with other observers. Rapid switching between two adaptive states is not limited to color vision. Similar rapid re-adaptation has been reported in sensorimotor paradigms, in which observers adapt to prisms that rotate or displace their visual field e.g. (Redding et al., 2005), or force fields that disturb their motor outcomes, e.g. (Wolpert & Flanagan, 2016). Visual mode switching also resembles context dependent learning that arises in conditioning and other memory paradigms. Mode switching appears to be a general solution to the problem of maintaining consistent behavior in a changing world.

## Materials and Methods

### Observers

Observers included author YL and 11 members (21 to 37 years of age) of the University of Minnesota community. All had normal color vision, as assessed by the Ishihara Color Blindness Test, and normal or corrected-to-normal (using contact lenses) visual acuity. None had worn red glasses for extended periods of time prior to this study. One of the observers recruited reported that she changed her criterion for unique yellow during the study, and her data showed very large variance in baseline across days. Her data were excluded from further analysis. Experimental procedures were approved by the University of Minnesota Institutional Review Board. All observers provided written, informed consent before the start of the study.

### Apparatus

Visual stimuli were presented on a NEC MultiSync FP2141 cathode ray tube monitor, with screen resolution of 1024*768 pixels, and a refresh rate of 85 Hz. The monitor was calibrated using a Photo Research PR655 spectroradiometer, with gun outputs linearized through look-up tables. Background luminance was 41.85 candela/m*2*. All visual stimuli were delivered in Matlab using the psychophysical toolbox (Brainard, 1997). Viewing distance was maintained at 50 cm with a chinrest.

### Glasses

Observers wore a commercial pair of bright red glasses made by *SomniLight (Shawnee, KS)*. Black baffling was added on the top of the frame to prevent light from bypassing the glasses from above. The glasses filter out most of the light at short wavelengths and let pass most of the light at long wavelengths. We measured the glasses transmittance by placing the glasses in front of the spectroradiometer and recording the light from sunlight. The spectral transmission of the glasses (Fig. 1A) shows that the transmittance is above 90% at wavelengths over 620 nm, and less than 10% at wavelengths below 550 nm.

To characterize the effect of the glasses on our testing display, we measured the gamut of the monitor with and without the glasses. Fig. 1B demonstrates that the gamut of the monitor seen through the glasses becomes compressed and shifts towards red chromaticity.

### Procedure

In the main experiment, observers wore the glasses for 5 one-hour periods per day, for 5 consecutive days. On each day, observers came to the lab in the morning and wore the red glasses for 1 hour, while participating in a testing session. Then, they left the lab and attended to their routine everyday activities. They were asked to put on the glasses again one hour after they took off the glasses in the lab. During the day, they wore the glasses for three one-hour periods, each separated by one hour without glasses. At the end of the fourth one-hour period without glasses, they came back to the lab for a second testing session, identical to that in the morning. Fig. 1C, upper panel, illustrates the procedure of the experiment. In a follow-up test session conducted about one month after the main experiment, observers came back and performed one additional and identical testing session.

During the test sessions, observers adjusted the color of a 0.5-degree square centered on a background image of a naturalistic environment (an office scene). A black square of 5.7 degrees separated the test patch from the background image (Fig. 1D). The goal was to set the small square to unique yellow. The small patch was presented for 200 ms at 1.5 sec intervals. To make adjustments observers pressed the left and down arrow buttons to reduce redness in the patch, right and up arrow buttons to reduce greenness in the patch, and then pressed the space bar when they had set the patch to appear neither reddish nor greenish. The left and right arrow buttons were for coarse adjustments, and the up and down arrow buttons were for finer adjustments. Observers had 20 sec at the most in each trial to make the adjustments.

Stimuli were created using a modified version of the MacLeod-Boynton color space (MacLeod & Boynton, 1979), scaled and shifted so that the origin corresponds to a nominal white point of Illuminant C and so that sensitivity is roughly equated along the LM and S axes (Webster et al., 2000):

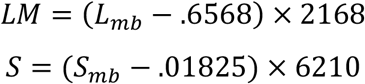

All settings fell along the nominally iso-luminant plane of this space when not wearing the glasses in order to reduce brightness effects on the judgements. The chromatic contrast was also kept constant during the adjustment procedure, thus results are shown in “Hue Angle,” where luminance and contrast were held constant. The stimuli were not adjusted for the glasses, and thus were likely not held at strictly constant luminance or contrast for judgements made while the glasses were on. Observers could adjust the color of the test patch with coarser or finer steps of 5 or 1 degree respectively per button press.

At the beginning of each test session, observers performed five one-minute blocks of this task with natural vision. Then, they put the glasses on and immediately did the task again. They were also tested after 10 min, 25 min, 40 min, and 55 min of wearing the glasses. Between tests observers took a short walk and watched videos of their choice on computer or phone. After 1 hour, observers removed the glasses and were immediately tested again. Further tests were performed 10 min, 20 min, and 30 min after removing the glasses. The full test procedure is illustrated in the lower panel of Fig. 1C. Critically, observers completed all tests in a fully lit room, with the aim of measuring perceptual experience in a context like their natural environment while adapting to the glasses.

### Relative cone absorption analysis

In order to compare unique yellow settings with and without the glasses, we also characterized the results in terms of relative absorptions of the cone photoreceptors computing using Stockman & Sharpe (2000) cone fundamentals (Stockman & Sharpe, 2000). We computed relative cone absorptions as follows: First, we calculated the spectra of the unique yellow settings by multiplying the RGB values of the observers’ settings by the gun spectra of the monitor and summing the outputs of the three guns. For the settings made with the glasses on, we further multiplied the monitor spectra by the transmission spectrum of the glasses. The spectra of the settings were then multiplied by the cone fundamentals to compute cone absorptions. Lastly, the absorptions were converted into relative cone absorptions (corresponding to the L vs. M cardinal axis in MacLeod–Boynton color space (MacLeod & Boynton, 1979)), computed by the ratio of L/(L+M), with and without the glasses correspondingly.

## Acknowledgements

This study was funded by NSF-BCS 1558308.

## References

Belmore, S. C., & Shevell, S. K. (2008). Very-long-term chromatic adaptation: test of gain theory and a new method. Visual Neuroscience, 25(3), 411–414. https://doi.org/10.1017/S0952523808080450

Belmore, S. C., & Shevell, S. K. (2011). Very-long-term and short-term chromatic adaptation: Are their influences cumulative? Vision Research, 51(3), 362–366. https://doi.org/10.1016/j.visres.2010.11.011

Boynton, R. M., & Whitten, D. N. (1970). Visual adaptation in monkey cones: recordings of late receptor potentials. Science (New York, N.Y.), 170(3965), 1423–1426. https://doi.org/10.1126/science.170.3965.1423

Brainard, D. H. (1997). The Psychophysics Toolbox. Spatial Vision, 10(4), 433–436. https://doi.org/10.1163/156856897X00357

Brainard, D. H., & Stockman, A. (2010). Colorimetry. In: Bass M, editor. OSA Handbook of Optics. 3rd. McGraw-Hill; New York. pp. 10.11–11.56

Brainard, D. H., & Radonjić, A. (2014). Color constancy. In The new visual neurosciences (eds. L.M. Chalupa & J.S. Werner), pp. 545–556. Cambridge, MA, MIT Press.

Clifford, C. W., Webster, M. A., Stanley, G. B., Stocker, A. A., Kohn, A., Sharpee, T. O., & Schwartz, O. (2007). Visual adaptation: Neural, psychological and computational aspects. Vision Research, 47(25), 3125–3131. https://doi.org/10.1016/j.visres.2007.08.023

de La Hire, P. (1694). Mémoires De Mathématique Et De Physique. De L’Imprimerie Royale.

Delahunt, P. B., & Brainard, D. H. (2004). Does human color constancy incorporate the statistical regularity of natural daylight? Journal of Vision, 4(2), 57–81. https://doi.org/10.1167/4.2.1

Dragoi, V., Sharma, J., Miller, E. K., & Sur, M. (2002). Dynamics of neuronal sensitivity in visual cortex and local feature discrimination. Nature Neuroscience, 5(9), 883–891. https://doi.org/10.1038/nn900

Eisner, A., & Enoch, J. M. (1982). Some effects of 1 week’s monocular exposure to long-wavelength stimuli. Perception & Psychophysics, 31(2), 169–174. https://doi.org/10.3758/BF03206217

Engel, S. A., & Furmanski, C. S. (2001). Selective adaptation to color contrast in human primary visual cortex. The Journal of Neuroscience: The Official Journal of the Society for Neuroscience, 21(11), 3949–3954. https://doi.org/10.1523/JNEUROSCI.21-11-03949.2001

Engel, S. A., Wilkins, A. J., Mand, S., Helwig, N. E., & Allen, P. M. (2016). Habitual wearers of colored lenses adapt more rapidly to the color changes the lenses produce. Vision Research, 125, 41–48. https://doi.org/10.1016/j.visres.2016.05.003

Eskew, R. T., & Richters, D. P. (2008). Potential mechanisms of long-term adaptation in color vision, and a failure to find evidence for them. Journal of Vision, 8(17), 26–26. https://doi.org/10.1167/8.17.26

Fairchild, M. D., & Lennie, P. (1992). Chromatic adaptation to natural and incandescent illuminants. Vision Research, 32(11), 2077–2085. https://doi.org/10.1016/0042-6989(92)90069-U

Fairchild, M. D., & Reniff, L. (1995). Time course of chromatic adaptation for color-appearance judgments. Journal of the Optical Society of America. A, Optics, image science, and vision, 12(5), 824–833. https://doi.org/10.1364/josaa.12.000824

Foster, D. H. (2011). Color constancy. Vision Research, 51(7), 674–700. https://doi.org/10.1016/j.visres.2010.09.006

Grzywacz, N., & de Juan, J. (2003). Sensory adaptation as Kalman filtering: Theory and illustration with contrast adaptation. Network: Computation in Neural Systems, 14(3), 465–482. https://doi.org/10.1088/0954-898X_14_3_305

Hill, A. R., & Stevenson, R. W. (1976). Long-term adaptation to ophthalmic tinted lenses. Modern Problems in Ophthalmology, 17, 264–272.

Jameson, D., & Hurvich, L. M. (1989). Essay concerning color constancy. Annual Review of Psychology, 40, 1–22. https://doi.org/10.1146/annurev.ps.40.020189.000245

Kohler, I. (1963). The formation and transformation of the perceptual world. Psychological Issues, 3(4, Monogr. No. 12), 1–173.

Kohn, A. (2007). Visual adaptation: Physiology, mechanisms, and functional benefits. Journal of Neurophysiology, 97(5), 3155–3164. https://doi.org/10.1152/jn.00086.2007

Kording, K. P., Tenenbaum, J. B., & Shadmehr, R. (2007). The dynamics of memory as a consequence of optimal adaptation to a changing body. Nature Neuroscience, 10(6), 779–786. https://doi.org/10.1038/nn1901

Krauskopf, J., & Karl, G. (1992). Color discrimination and adaptation. Vision Research, 32(11), 2165–2175. https://doi.org/10.1016/0042-6989(92)90077-V

Krekelberg, B., van Wezel, R. J. A., & Albright, T. D. (2006). Adaptation in Macaque MT Reduces Perceived Speed and Improves Speed Discrimination. Journal of Neurophysiology, 95(1), 255–270. https://doi.org/10.1152/jn.00750.2005

Lee, B. B., Dacey, D. M., Smith, V. C., & Pokorny, J. (1999). Horizontal cells reveal cone type-specific adaptation in primate retina. Proceedings of the National Academy of Sciences of the United States of America, 96(25), 14611–14616. https://doi.org/10.1073/pnas.96.25.14611

MacLeod, D. I. A., & Boynton, R. M. (1979). Chromaticity diagram showing cone excitation by stimuli of equal luminance. Journal of the Optical Society of America, 69(8), 1183–1186. https://doi.org/10.1364/JOSA.69.001183

McDermott, K. C., Malkoc, G., Mulligan, J. B., & Webster, M. A. (2010). Adaptation and visual salience. Journal of Vision, 10(13), 17. https://doi.org/10.1167/10.13.17

Müller, J. R., Metha, A. B., Krauskopf, J., & Lennie, P. (1999). Rapid adaptation in visual cortex to the structure of images. Science, 285(5432), 1405–1408. https://doi.org/10.1126/science.285.5432.1405

Murch, G. M., & Hirsch, J. (1972). The McCollough effect created by complementary afterimages. The American Journal of Psychology, 85(2), 241–247. https://doi.org/10.2307/1420664

Neitz, J., Carroll, J., Yamauchi, Y., Neitz, M., & Williams, D. R. (2002). Color perception is mediated by a plastic neural mechanism that is adjustable in adults. Neuron, 35(4), 783–792. https://doi.org/10.1016/S0896-6273(02)00818-8

Pearce, B., Crichton, S., Mackiewicz, M., Finlayson, G. D., & Hurlbert, A. (2014). Chromatic illumination discrimination ability reveals that human colour constancy is optimised for blue daylight illuminations. PloS one, 9(2), e87989. https://doi.org/10.1371/journal.pone.0087989

Radonjić, A., & Brainard, D. H. (2016). The nature of instructional effects in color constancy. Journal of Experimental Psychology: Human Perception and Performance, 42(6), 847–865. https://doi.org/10.1037/xhp0000184

Redding, G. M., Rossetti, Y., & Wallace, B. (2005). Applications of prism adaptation: A tutorial in theory and method. Neuroscience & Biobehavioral Reviews, 29(3), 431–444. https://doi.org/10.1016/j.neubiorev.2004.12.004

Rieke, F., & Rudd, M. E. (2009). The challenges natural images pose for visual adaptation. Neuron, 64(5), 605–616. https://doi.org/10.1016/j.neuron.2009.11.028

Rinner, O., & Gegenfurtner, K. R. (2000). Time course of chromatic adaptation for color appearance and discrimination. Vision Research, 40(14), 1813–1826. https://doi.org/10.1016/S0042-6989(00)00050-X

Rüttiger, L., Braun, D. I., Gegenfurtner, K. R., Petersen, D., Schönle, P., & Sharpe, L. T. (1999). Selective color constancy deficits after circumscribed unilateral brain lesions. The Journal of Neuroscience: The Official Journal of the Society for Neuroscience, 19(8), 3094–3106. https://doi.org/10.1523/JNEUROSCI.19-08-03094.1999

Seriès, P., Stocker, A. A., & Simoncelli, E. P. (2009). Is the homunculus “aware” of sensory adaptation? Neural Computation, 21(12), 3271–3304. https://doi.org/10.1162/neco.2009.09-08-869

Sharpee, T. O., Sugihara, H., Kurgansky, A. V., Rebrik, S. P., Stryker, M. P., & Miller, K. D. (2006). Adaptive filtering enhances information transmission in visual cortex. Nature, 439(7079), 936–942. https://doi.org/10.1038/nature04519

Sheth, B. R., & Shimojo, S. (2008). Adapting to an aftereffect. Journal of Vision, 8(3), 1–10. https://doi.org/10.1167/8.3.29

Smithson, H., & Zaidi, Q. (2004). Colour constancy in context: Roles for local adaptation and levels of reference. Journal of Vision, 4(9), 693–710. https://doi.org/10.1167/4.9.3

Stockman, A., & Sharpe, L. T. (2000). The spectral sensitivities of the middle-and long-wavelength-sensitive cones derived from measurements in observers of known genotype. Vision Research, 40(13), 1711–1737. https://doi.org/10.1016/S0042-6989(00)00021-3

Sugita, Y. (2004). Experience in early infancy is indispensable for color perception. Current Biology: CB, 14(14), 1267–1271. https://doi.org/10.1016/j.cub.2004.07.020

Tregillus, K. E. M., Werner, J. S., & Webster, M. A. (2016). Adjusting to a sudden “aging” of the lens. Journal of the Optical Society of America A, Optics, image science, and vision, 33(3), A129–A136. https://doi.org/10.1364/JOSAA.33.00A129

van Lier, R., Vergeer, M., & Anstis, S. (2009). Filling-in afterimage colors between the lines. Current Biology: CB, 19(8), R323–R324. https://doi.org/10.1016/j.cub.2009.03.010

Vinas, M., Sawides, L., de Gracia, P., & Marcos, S. (2012). Perceptual adaptation to the correction of natural astigmatism. PLoS one, 7(9), e46361. https://doi.org/10.1371/journal.pone.0046361

von Helmholtz, H. (1924). Helmholtz’s Treatise on Physiological Optics. (Trans. from the 3rd German ed.) (J. P. C. Southall, Ed.). Optical Society of America. https://doi.org/10.1037/13536-000

von Kries, J. (1902). Theoretische Studien über die Umstimmung des Sehorgans. In Festschrift der Albrecht-Ludwigs-Universität (pp. 143–158). Freiburg.

Wainwright, M. J. (1999). Visual adaptation as optimal information transmission. Vision Research, 39(23), 3960–3974. https://doi.org/10.1016/S0042-6989(99)00101-7

Wark, B., Fairhall, A., & Rieke, F. (2009). Timescales of inference in visual adaptation. Neuron, 61(5), 750–761. https://doi.org/10.1016/j.neuron.2009.01.019

Wark, B., Lundstrom, B. N., & Fairhall, A. (2007). Sensory adaptation. Current Opinion in Neurobiology, 17(4), 423–429. https://doi.org/10.1016/j.conb.2007.07.001

Webster, M. A. (2015). Visual Adaptation. Annual Review of Vision Science, 1, 547–567. https://doi.org/10.1146/annurev-vision-082114-035509

Webster, M. A., Miyahara, E., Malkoc, G., & Raker, V. E. (2000). Variations in normal color vision. II. Unique hues. Journal of the Optical Society of America. A, Optics, image science, and vision, 17(9), 1545–1555. https://doi.org/10.1364/josaa.17.001545

Webster, M. A., & Mollon, J. D. (1995). Colour constancy influenced by contrast adaptation. Nature, 373(6516), 694–698. https://doi.org/10.1038/373694a0

Weiss, D., Witzel, C., & Gegenfurtner, K. (2017). Determinants of Colour Constancy and the Blue Bias. i-Perception, 8(6), 2041669517739635. https://doi.org/10.1177/2041669517739635

Wissig, S. C., Patterson, C. A., & Kohn, A. (2013). Adaptation improves performance on a visual search task. Journal of Vision, 13(2), Article 6. https://doi.org/10.1167/13.2.6

Witzel, C., & Gegenfurtner, K. R. (2018). Color perception: Objects, constancy, and categories. Annual Review of Vision Science, 4(1), 475–499. https://doi.org/10.1146/annurev-vision-091517-034231

Wolpert, D. M., & Flanagan, J. R. (2016). Computations underlying sensorimotor learning. Current Opinion in Neurobiology, 37, 7–11. https://doi.org/10.1016/j.conb.2015.12.003

Wright, W. D., & Parsons, J. H. (1934). The measurement and analysis of colour adaptation phenomena. Proceedings of the Royal Society of London. Series B, Containing Papers of a Biological Character, 115(791), 49–87. https://doi.org/10.1098/rspb.1934.0029

Yang, J., Kanazawa, S., Yamaguchi, M. K., & Motoyoshi, I. (2015). Pre-constancy Vision in Infants. Current Biology: CB, 25(24), 3209–3212. https://doi.org/10.1016/j.cub.2015.10.053

Yehezkel, O., Sagi, D., Sterkin, A., Belkin, M., & Polat, U. (2010). Learning to adapt: Dynamics of readaptation to geometrical distortions. Vision Research, 50(16), 1550–1558. https://doi.org/10.1016/j.visres.2010.05.014

