## Supplemental Figures for "Visual mode switching learned through experience"

### Supplementary materials

#### Figures

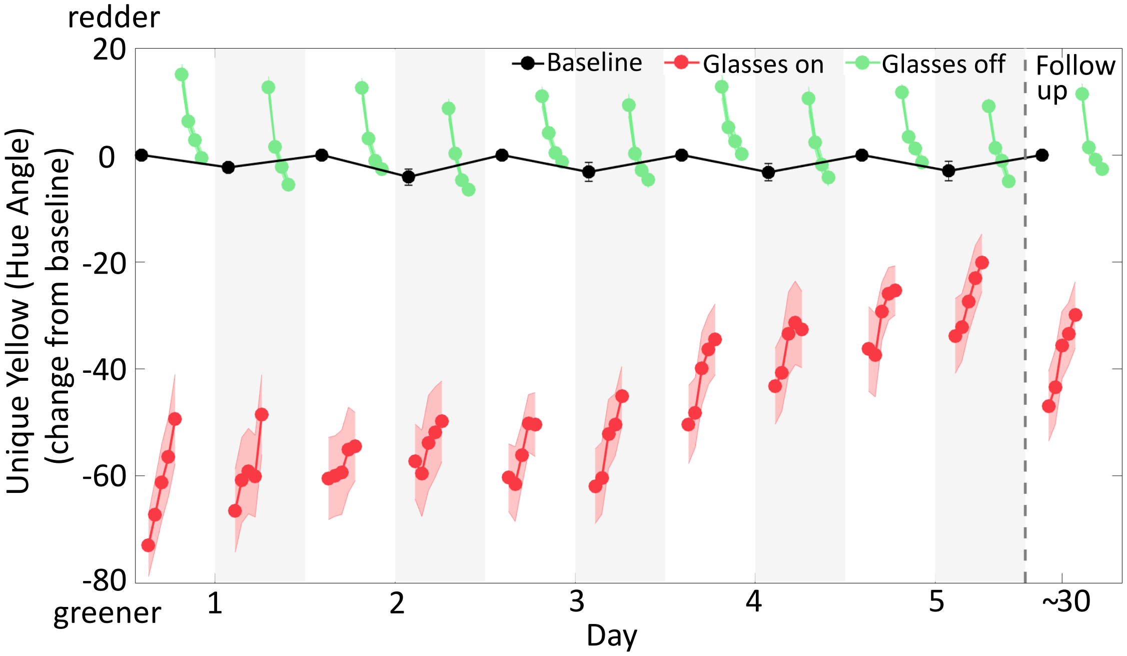


Fig. S1 Baseline-corrected results of the main experiment and the follow-up tests. Mean unique yellow settings represented in hue angle are plotted as a function of time for 5 days and the follow-up test. Data were normalized by subtracting the baseline value for each session, taken immediately before putting the glasses on. The black dots are baseline settings, made at the beginning of each test session with glasses off. The white background indicates morning sessions, and the light gray background indicates afternoon. The red dots plot settings with glasses on and the green dots are settings after removing the glasses. Successive symbols are plotted for each 5 min test. The red and green bands represent standard errors of the mean, computed across participants (N = 11). These baseline-corrected results showed an identical overall pattern across days as the uncorrected data, though some became slightly larger.

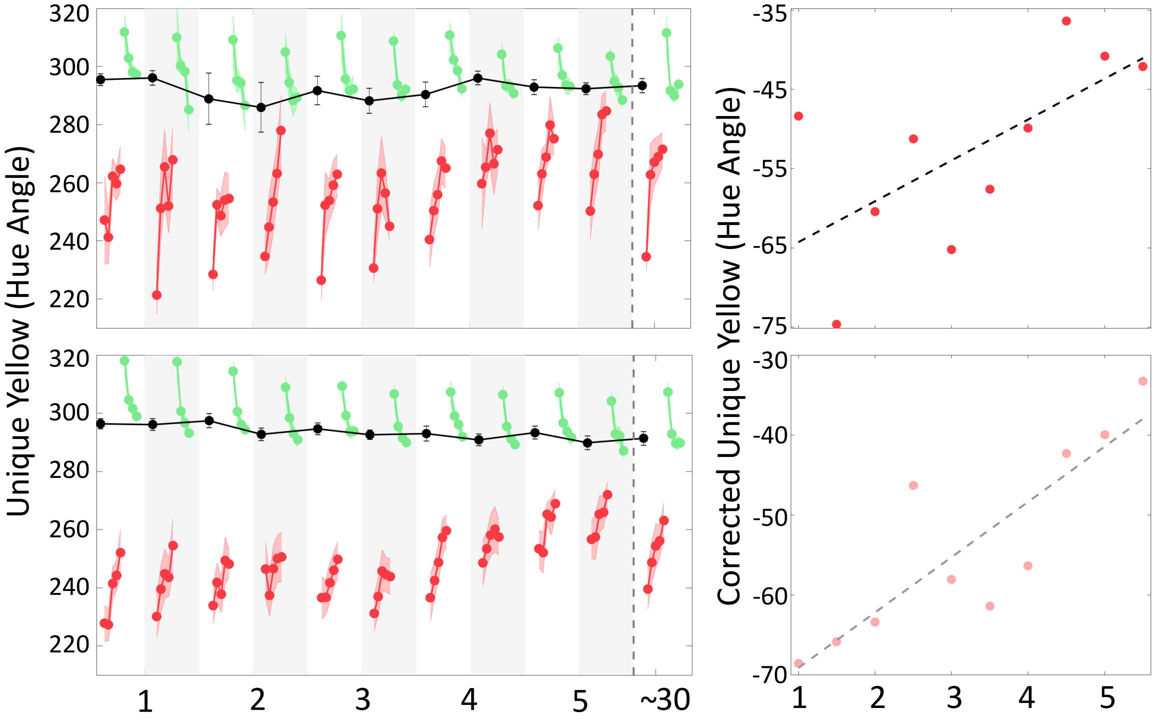


Fig. S2 Results of the first setting and the first block in the main experiment and the follow-up tests. The upper panel plots the very first unique yellow settings in each session, which were done within 20 seconds. Red dots in the scatter plot on the right show rapid effects of adaptation, which are the first settings with the glasses on. The black dashed line is the least-square fit; the rapid effects shifted significantly across days ($p = 0.03$). The lower panel plots the mean unique yellow settings within the first block in each session, during one-minute test. Pink dots in the scatter plot show rapid effects of adaptation, which are mean settings from the first 1 min test of each session with the glasses on. The gray dashed line is the least-square fit. The rapid effects shifted significantly across day ($p < 0.001$).
